# A single NPFR neuropeptide F receptor neuron that regulates thirst behaviors in Drosophila

**DOI:** 10.1101/2025.07.03.662850

**Authors:** Donnoban Orozco Ramirez, Brian P. Wang, Dan Landayan, Gagandeep Kaur, Fred W. Wolf

## Abstract

Thirst is a strongly motivated internal state that is represented in central brain circuits that are only partially understood. Water seeking is a discrete step of the thirst behavioral sequence that is amenable to uncovering the mechanisms for motivational properties such as goal-oriented behavior, value encoding, and behavioral competition. In Drosophila water seeking is regulated by the NPY-like neuropeptide NPF, however the circuitry for NPF-dependent water seeking is unknown. To uncover the downstream circuitry, we identified the NPF receptor NPFR and the neurons it is expressed in as being acutely critical for thirsty water seeking. Refinement of the NPFR pattern uncovered a role for a single neuron, the L1-l, in promoting thirsty water seeking. The L1-l neuron increases its activity in thirsty flies and is involved in the regulation of dopaminergic neurons in long-term memory formation. Thus, NPFR and its ligand NPF, already known for its role in feeding behavior, are also important for a second ingestive behavior.

**Significance Statement:** Understanding how a single motivated behavior is represented in the neural circuitry of the brain will help uncover drive-specific and universal encoding mechanisms for motivation. Thirst is useful because it is relatively simple compared to feeding behavior and it is strongly motivated. The fly Drosophila, with its straightforward genetics and complete connectome, is helpful in determining a complete circuit for thirsty water seeking. Here we discover a single neuron in the fly brain, the L1-l, that actively receives input from the NPY-like neuropeptide NPF to promote water seeking. Prior findings suggest that the L1-l modulates valence inputs into sensory processing centers, suggesting a similar function in thirsty seeking.

**Visual Abstract:** 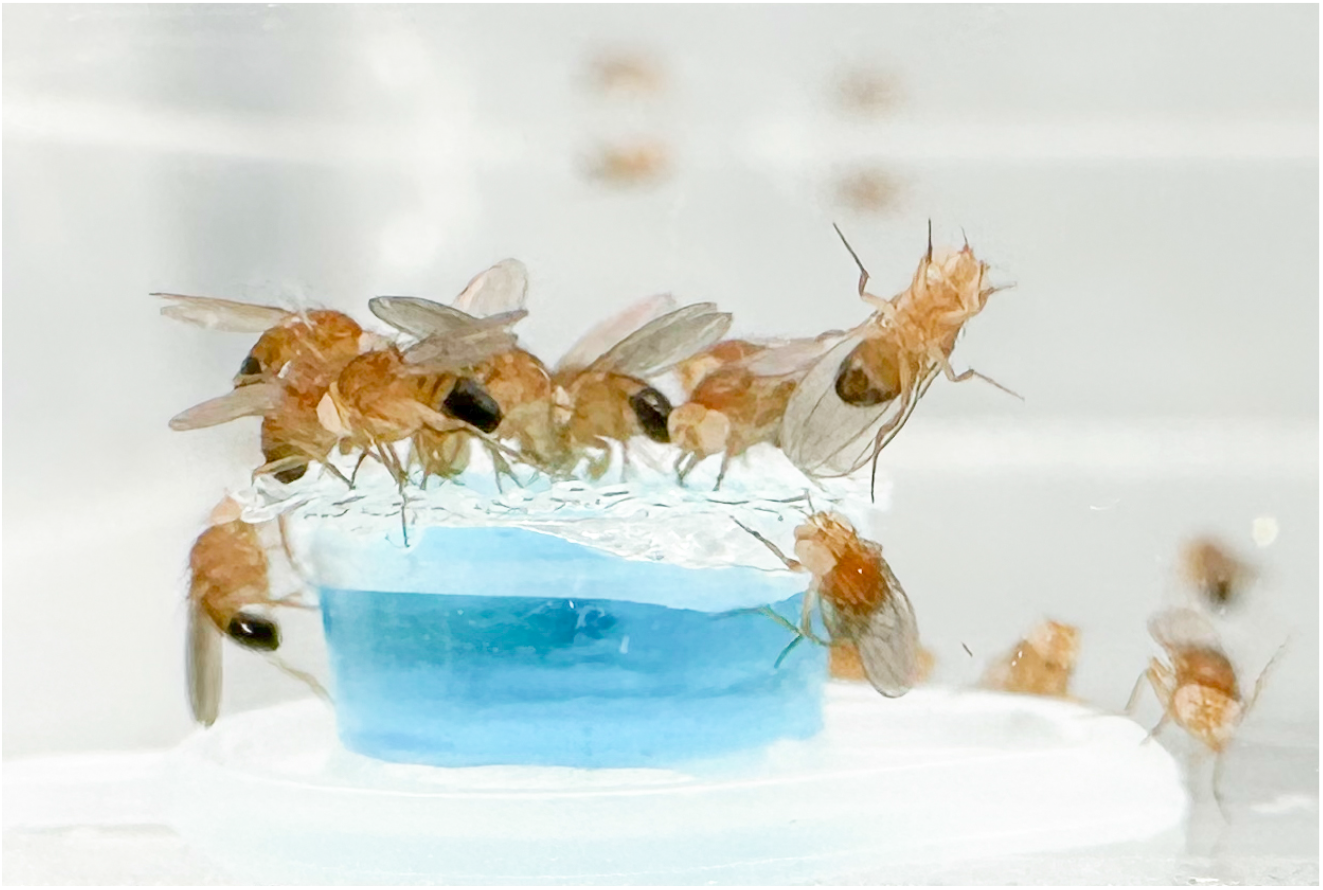

## Introduction

Thirst is an internal state that drives the seeking and consumption of water to restore osmotic homeostasis (Fitzsimons, 1972). The representation of thirst in the brain provides a means for understanding the neural encoding of motivational states, competition between conflicting needs, and behavioral sequencing. Uncovering the neural circuitry that supports specific steps of thirst behaviors will help reveal the bases for some of these complex computations. Animals become thirsty in a graded manner as their interstitial fluids increase in osmolarity. Thirst initiates a set of behaviors that often occur in sequence: searching for a source of water, determining if the water is palatable, water ingestion, and termination of water ingestion at satiety. It is accompanied by changes in sensory acuity, an escalating negative valence state and shorter duration positive valence anticipatory states, suppression of competing behaviors, and neuroendocrine signaling to minimize water loss (Betley et al., 2015; Gizowski et al., 2016; Gizowski and Bourque, 2018; Gáliková et al., 2018; Zandawala et al., 2021; Chu et al., 2024; Encarnacion-Rivera et al., 2025).

Thirsty Drosophila exhibit behaviors similar to other thirsty animals. Water deprivation changes humidity aversion to attraction, promotes directed flight and locomotion up humidity gradients, and increases water intake (Dethier and Evans, 1961; Ji and Zhu, 2015; Limbania et al., 2023). Osmosensory neurons detect osmotic changes in the circulating hemolymph and direct water ingestion through a simple circuit to neuroendocrine neurons in the pars intercerebralis (Jourjine, 2017; González Segarra et al., 2023). Humidity is detected by receptors on the antennae and humidity information is transmitted to a broad array of higher order central brain circuits (Enjin et al., 2016; Frank et al., 2017; Knecht et al., 2017; Marin et al., 2020; Chu et al., 2024). Thirst-dependent water seeking and humidity preference are controlled by central brain circuitry that includes neurons in the mushroom body learning and memory center, dopaminergic neurons that project to the mushroom body and provide valuation information, and a projection neuron that connects the primarily sensory subesophageal zone (SEZ) to the superior medial protocerebrum (SMP) (Lin et al., 2014; Ji and Zhu, 2015; Landayan et al., 2021). The projection neuron, Janu-AstA, releases the Galanin/Kisspeptin/Spexin neuropeptide ortholog Allatostatin A (AstA) onto neurons that release the Neuropeptide Y (NPY) ortholog Neuropeptide F (NPF) to regulate thirsty water seeking.

These findings led us to search for the next downstream layer of the thirsty water seeking circuit by characterizing the role of NPFR, the only known receptor for NPF. NPFR regulates many processes in flies including food intake, energy homeostasis, the effect of temperature on circadian rhythms, courtship, stress, sensitivity to ethanol, and learning and memory (Wen et al., 2005; Wu et al., 2005; Krashes et al., 2009; Xu et al., 2010; Feng et al., 2021; Ryvkin et al., 2021, 2024; Yoshinari et al., 2021; Gao et al., 2024; Yuan et al., 2024). The interrelationship of these behaviors, if any, remains unclear. Assignment of specific functions of NPFR to specific neurons or cells can help untangle potential relationships and neural functions. For example, consolidation of long-term memory requires NPFR in the aversive-encoding PPL1 dopamine neurons, whereas circadian-dependent locomotor activity in the late afternoon is suppressed by NPFR in LPN neurons when it is cold, indicating separate functions of NPFR in these complex behaviors (Feng et al., 2021; Yuan et al., 2024). Here, we discovered that NPFR functions in a single neuron, the L1-l, to promote thirsty water seeking.

## Methods

### Genetics and culturing conditions

All strains were outcrossed for at least five generations to the *Berlin* strain of *Drosophila melanogaster* that contained the *w*^*1118*^ mutation; this strain is indicated as ‘control’ in the figures. Crosses were incubated at 25°C and 60% humidity on a 12 hr light, 12 hr dark schedule. Adult flies were removed from crosses after 2-3 days to create age-matched cohorts of progeny. Progeny were collected using CO_2_ anesthesia 2-4 days after eclosion and then held 1-2 days prior to testing. Male flies were used for all experiments. Strains and sources are listed in **Table I**.

**Table I.**
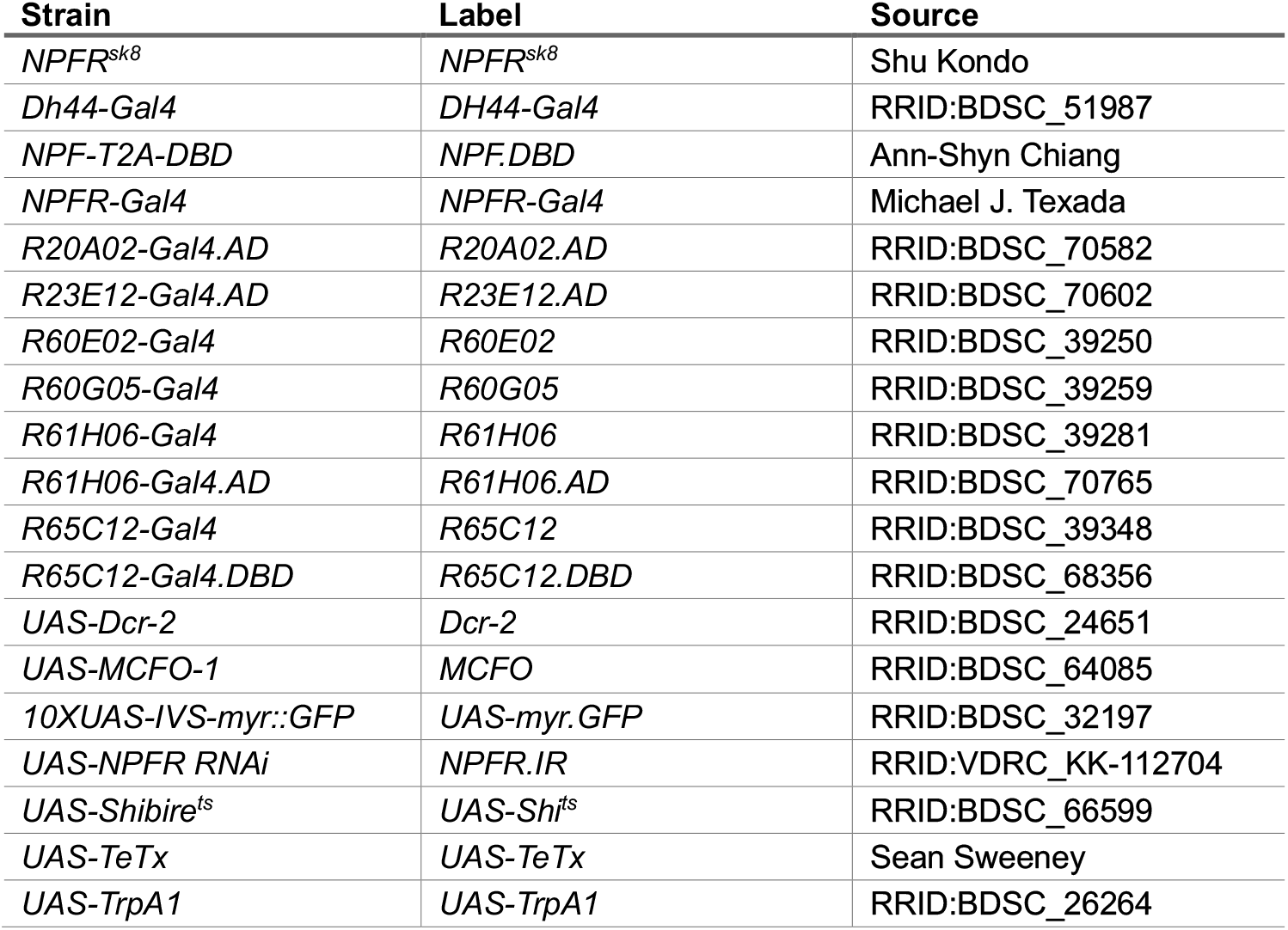
Strains.

### Water seeking and foraging behaviors

Flies were deprived of water in culturing vials lined with a 2.5 × 15 cm strip of Whatman 3 MM filter paper that was infused with 5% sucrose and then dried. During the water deprivation period the flies were held in an incubator at 25°C and 60% humidity. Thin-walled Plexiglas behavioral chambers (IO Rodeo, Pasadena CA) were designed with two side-by-side arenas, each arena measuring 45 × 75 × 10 mm. The chamber and water source were acclimated to the testing temperature in a Peltier incubator (IN45, Torrey Pines Scientific). Flies were acclimated in the chambers for 10 min prior to introducing the water source. Humidity was monitored (EK-H4 multiplexer with SHT71 sensors, Sensirion) and maintained at 30–40% relative humidity. Flies were filmed from above with a Logitech C510 web camera at 30 fps with the arena placed on white light LED panel (Edmund Optics). Sources of water (containing 0.2% w/v erioglaucine, 229730050, Fisher Scientific) were presented to flies in an overturned cap of a 0.65 mL microcentrifuge tube. Water was made inaccessible by glueing (Cat No. 698, DAP) a 300 μm mesh grid (U-CMN-300-A, Components Supply) onto the cap. The number of flies on the inaccessible water source was divided by the total number of flies in the arena to determine the fraction of flies on the source. Food foraging behavior followed the same protocol as for water seeking, except that flies were food deprived in culturing vials lined with Whatman 3MM soaked in water, and food deprived flies were placed in the open field arena and presented with a small chunk of cane sugar (La Perruche Pure Cane Rough Cut Cubes) resting in an overturned 1.5 mL microcentrifuge tube cap.

### Water intake

Flies were glued to microscope slides with clear nail polish while cold anesthetized and allowed to recover at 25°C and 60% humidity for 1-2 hours. Consumption time was measured by repeatedly presenting the proboscis with a water drop extruded from the end of a 5 μL calibrated micropipette (53432-706, VWR), until flies failed to extend the proboscis to four sequential presentations. 0.2% w/v erioglaucine was added to help with quantification. Ingestion behavior was recorded on a stereoscope with a AmScope MD500 microscope eyepiece camera at 30 fps at 640 x 480. Pumping rate was determined with manual counts of video recordings of pharyngeal contractions during the initial 10 s of the first engagement, or shorter if the flies disengaged earlier.

### Immunohistochemistry

Adult brains were dissected in 1x phosphate buffered saline (PBS) (BP399, Fisher Scientific) containing 0.05% Triton-X 100 (X100-500ML, Sigma Aldrich) (0.05% PBT), fixed overnight at 4°C in 0.05% PBT with 2% paraformaldehyde (50-980-493, Fischer Scientific). Brains were then washed with 0.1% PBT 5 times for 20 minutes at room temperature, blocked in 0.5% PBT with 5% normal goat serum (733250, Lampire Biological Laboratories) and 0.5% bovine serum albumin (A7906-50G, Sigma-Aldrich) for 2 hours at room temperature, and incubated with primary antibody diluted in block for 1-3 days at 4°C. Brains were washed with 0.1% PBT for 5x 20 minutes at room temperature, incubated in secondary antibody diluted in 0.1% PBT for 1-2 days at 4°C, washed with 0.1% PBT for 5x 20 minutes at room temperature, and equilibrated into Vectashield (H-1000, Vector Laboratories) overnight at 4°C. Samples were mounted on slides and imaged on a Zeiss LSM-880 confocal microscope with a 20x objective. Image stacks were processed in Fiji, and brightness and contrast were adjusted in Adobe Photoshop. For MultiColor FlpOut experiments, flies were heat shocked in a 37°C water bath for 40-45 minutes before resting at 25°C and 60% humidity for 2 days, and then dissected and stained as above. Antibodies used were rabbit anti-GFP, 1:1000 (A6455, Invitrogen/Fisher), chicken anti-GFP, 1:1000 (AB13970, Abcam) mouse anti-Bruchpilot, 1:25 (NC82, Developmental Studies Hybridoma Bank), rabbit anti-NPFR, N-terminal, 1:25 (RB-19-0003-100, RayBiotech), goat anti-rabbit Alexa Fluor 488, 1:350 (A11034, Molecular Probes), goat anti-mouse Alexa Fluor 594, 1:350 (A11032, Molecular Probes), donkey anti-chicken FITC, 1:350 (SA172000, Pierce), and goat anti-rabbit Alexa Fluor 594, 1:350 (A11037, Molecular Probes).

### Experimental design and statistical analysis

Groups of 20 genetically identical flies of the same age constituted an *n* = 1 for all behavioral assays, except for water intake where individual animals were assessed. No statistical methods were used to predetermine sample sizes. An *n* = 8 was previously determined to give discriminative power for the water seeking and food foraging open field assays (Landayan et al., 2021). For water intake an *n* = 20-30 individuals follows the design in prior reports (Jourjine, 2017). Experiments were typically performed with flies from 2-3 separate crosses and across 2-3 days. Statistical analysis was carried out in Prism v10.5.0. All statistical tests and outcomes are detailed in **Table II**. Dots overlaid on bar graphs are the value measured for individual groups, or for individuals for water intake. No data was excluded using statistical methods. Error bars are the standard error of the mean, except for water intake, where data is represented as box and whiskers that represent the minimum and maximum values.

**Table II.**
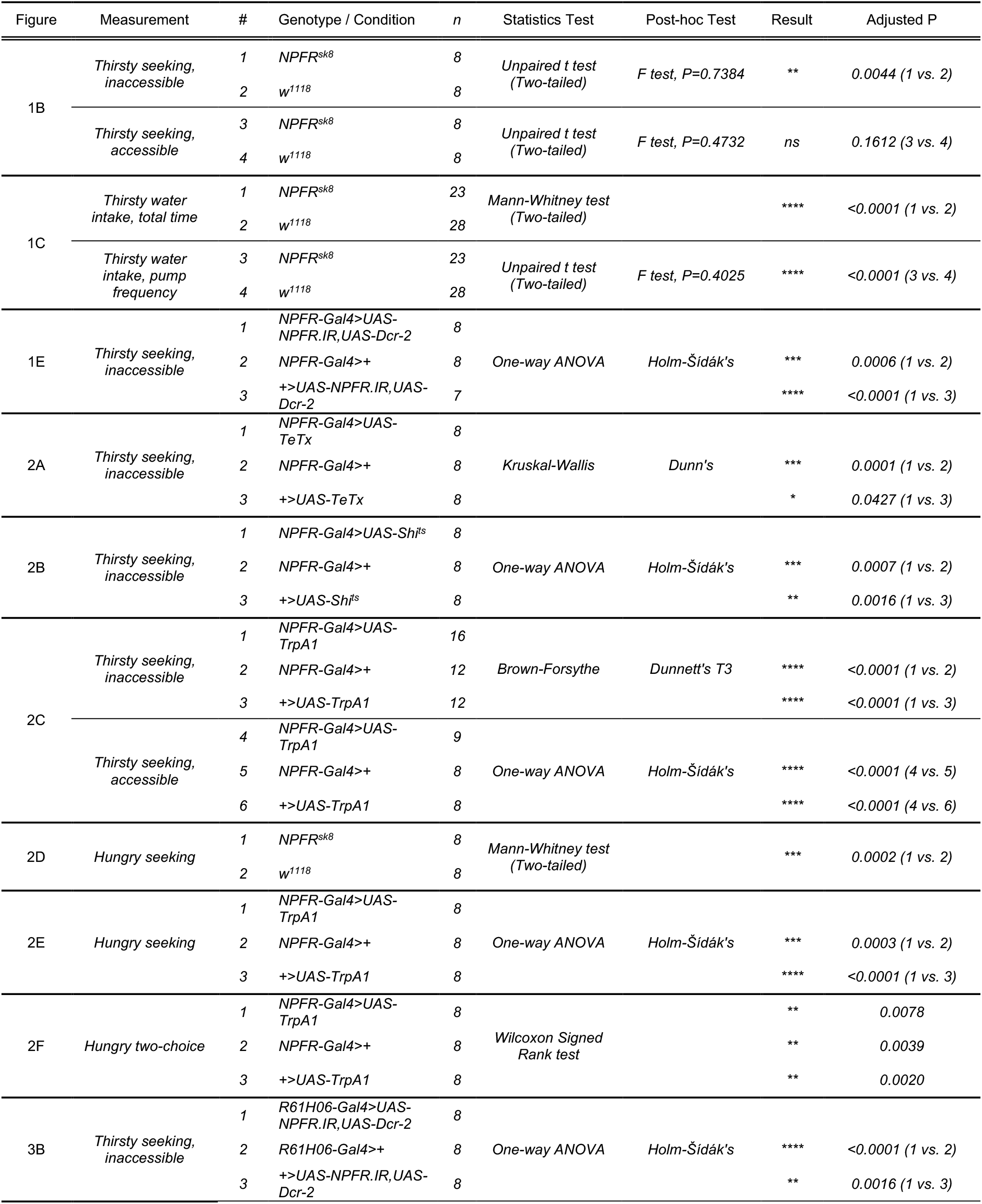
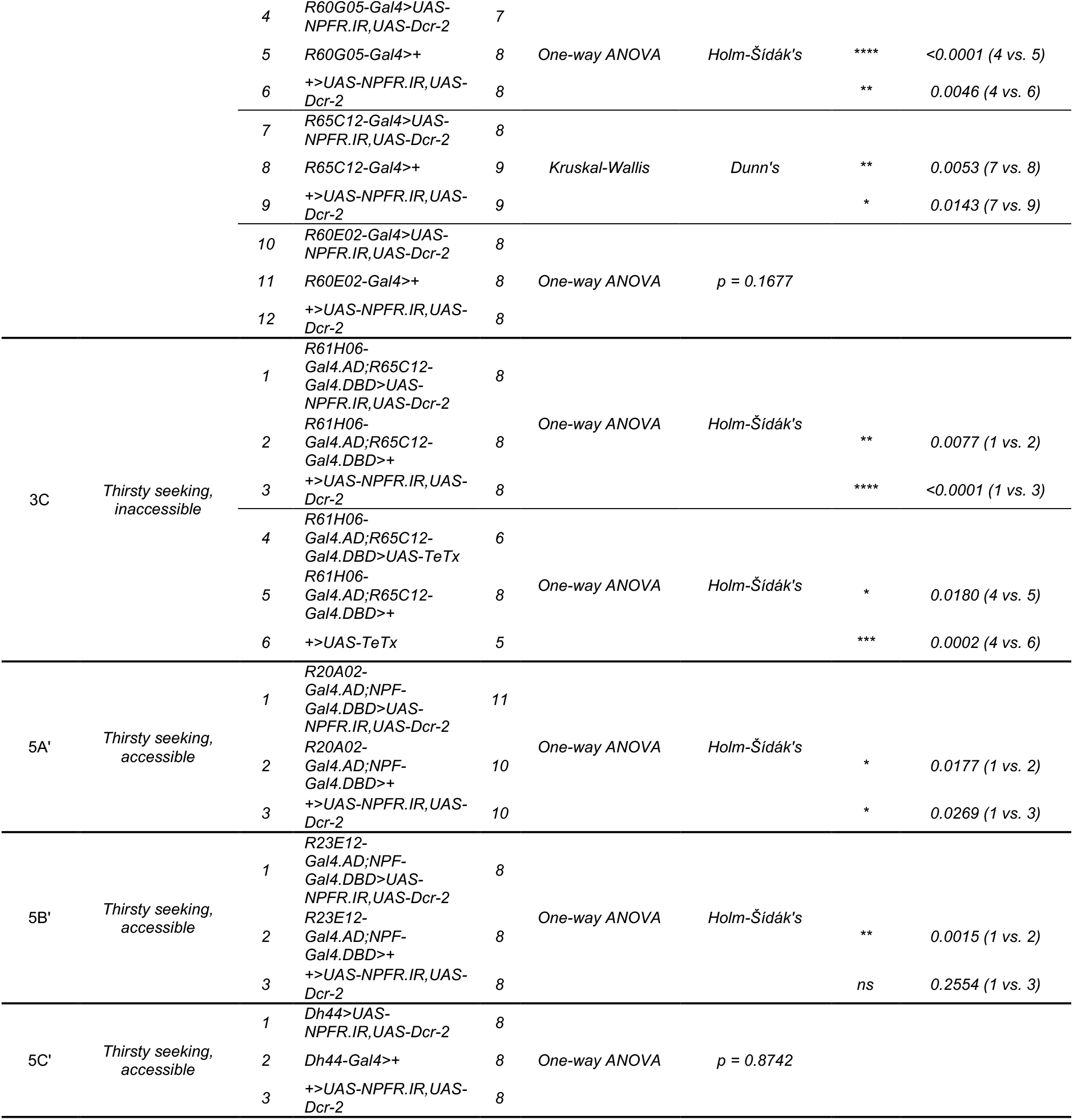
Genotypes and statistical analyses.

## Results

### NPFR acts in NPFR-Gal4 neurons to promote thirsty water seeking and intake

To ask if NPFR functions in thirst, we tested a loss-of-function *NPFR* mutant for thirsty water seeking. *NPFR*^*sk8*^ is a frameshift mutation near the 5’ end of the *NPFR* open reading frame that eliminates NPFR expression (Ameku et al., 2018). We measured water seeking in an open field assay, where a group of 20 flies are placed in an arena with a small reservoir of water that is made inaccessible by placing a mesh grid over the reservoir (**Figure 1A**). The reservoir creates a shallow humidity gradient, and thirsty flies travel up the gradient to occupy the inaccessible water source in a water deprivation- and time-dependent manner. *NPFR* mutant flies showed reduced water seeking when thirsty (**Figure 1B**). By contrast, *NPFR* mutants performed normally when given an open, accessible water source, indicating that *NPFR* mutants do not have general behavioral deficits (**Figure 1B**). To determine if NPFR regulates thirst broadly or is specific to the water seeking step, we tested water ingestion by presenting flies with a small drop of water on their proboscis, measuring ingestion time and pharyngeal pumping frequency. Thirsty *NPFR* mutants drank for a longer time and had a decreased pumping frequency (**Figure 1C**). Thus, flies lacking NPFR have blunted water seeking, and apparent compensatory changes in water intake time and rate. To identify the site of NPFR action, we asked if *NPFR* acts in neurons labeled by an *NPFR-Gal4* transgene. In this transgene *Gal4* is inserted immediately 3’ to the NPFR initiator methionine in an 18 kb genomic clone of the *NPFR* locus that is integrated into the genome at the *attP40* landing site (**Figure 1D**). *NPFR-Gal4* driving expression of myristoylated GFP (*UAS-myr*.*GFP*) revealed expression in approximately 100 neurons. The pattern included the NPF-positive neurons L1-l and P1, neuroendocrine cells in the pars intercerebralis, additional unidentified central brain neurons, and about 20 neurons in the ventral nerve cord (**Figure 1D**). When *NPFR* RNAi (*UAS-NPFR*.*IR*) was driven by *NPFR-Gal4*, thirsty flies showed reduced water seeking (**Figure 1E**). Thus, NPFR acts in neurons in the *NPFR-Gal4* pattern to promote thirsty water seeking.

**Figure 1.**
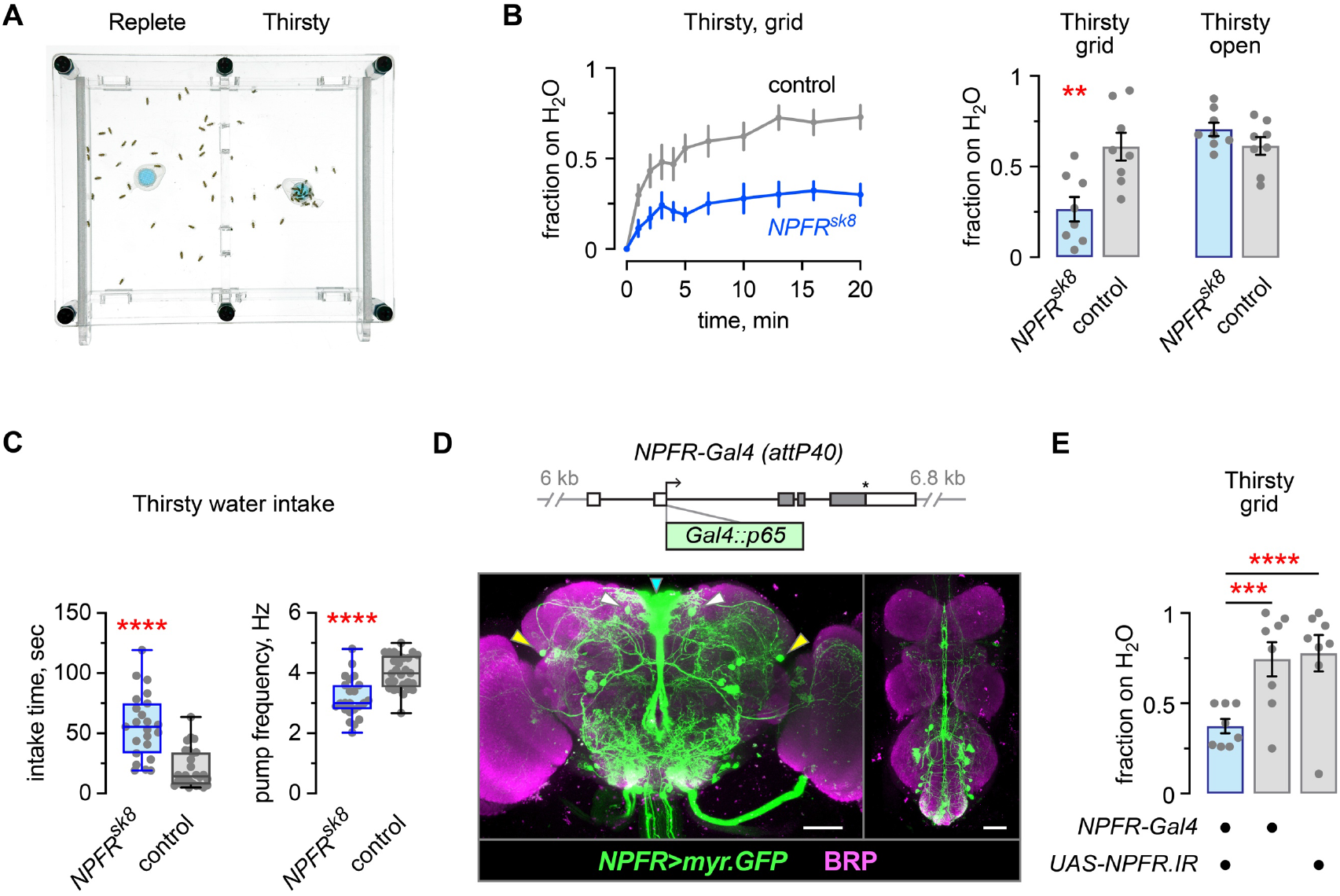
NPFR regulates thirst behaviors in *NPFR-Gal4* neurons. **A**. Open field device for assessing water seeking, with an inaccessible water source placed at the center of each arena. **B**. Left: Time course of flies occupying a gridded inaccessible water source over time, genetic control flies and *NPFR*^*sk8*^ strong loss-of-function mutants. Flies were water deprived on dry sucrose for 16 hours at 25°C and 60% humidity. Right: *NPFR*^*sk8*^ mutants exhibit decreased thirsty water seeking specifically to an inaccessible source. Left: unpaired t test, two-tailed, *p* = 0.0044; right: Unpaired t test, two-tailed, *p* = 0.1612. **C**. Increased intake time and decreased pharyngeal pumping rate in *NPFR*^*sk8*^ mutants. Left: Mann-Whitney test, two-tailed, *p* < 0.0001; right: Unpaired t test, two-tailed, *p* < 0.0001. **D**. Genomic structure (top) and expression (bottom) of *NPFR-Gal4* in the Drosophila central brain (left) and ventral nerve cord (right), immunostained for GFP and the presynaptic protein Bruchpilot (BRP) to reveal the synaptic neuropil. Arrowheads point to the L1-l (yellow), P1 (white), and pars intercerebralis (cyan) neurons. Scale bars: 50μm. **E**. *NPFR RNAi* (*UAS-NPFR*.*IR*,*UAS-Dcr-2*) in *NPFR-Gal4* reduced thirsty water seeking. One-way ANOVA with Holm-Šídák’s multiple comparisons test; *Gal4/UAS* vs. *Gal4, p* = 0.0006; *Gal4/UAS* vs. *UAS, p* < 0.0001.

### Activity of NPFR-Gal4 neurons is acutely required for thirsty water seeking and for food foraging

We next determined the role of neuronal activity in the *NPFR-Gal4* pattern. We inactivated the neurons in two ways. First, expression of the tetanus toxin light chain (*UAS-TeTx*) that blocks presynaptic release of synaptic vesicles in *NPFR-Gal4* reduced thirsty water seeking (**Figure 2A**). Second, acute and rapid inactivation of NPFR neurons using a temperature sensitive dominant negative Shibire (dynamin, *UAS-Shi*^*ts*^) also reduced thirsty water seeking (**Figure 2B**). Thus, NPFR neurons promote thirsty water seeking during the water seeking task. Finally, to test if NPFR neurons are sufficient to promote water seeking, we acutely and rapidly activated them using the TrpA1 cation channel (*UAS-TrpA1*) in water replete flies. NPFR neuron activation increased water seeking behavior (**Figure 2C**). Thus, NPFR neurons are both necessary and sufficient for water seeking behavior, and they regulate water seeking via the NPFR receptor. NPF regulates both feeding and thirst-driven behavior (Beshel and Zhong, 2013; Landayan et al., 2021). We found that hungry *NPFR* mutants had a strong deficit in foraging for dry sucrose (**Figure 2D**). Acute TrpA1 activation of NPFR neurons also strongly reduced feeding behavior (**Figure 2E**). Since activation of NPFR neurons promoted water seeking but inhibited feeding behavior, we predicted that hungry flies would prefer water seeking when NPFR neurons are activated. Hungry but water-replete *NPFR>TrpA1* flies given a choice between water and dry sucrose preferred the water source (**Figure 2F**). Thus, ongoing activity of the NPFR neurons promotes water seeking and inhibits feeding behavior, even when water seeking is mismatched to the prevailing internal state.

**Figure 2.**
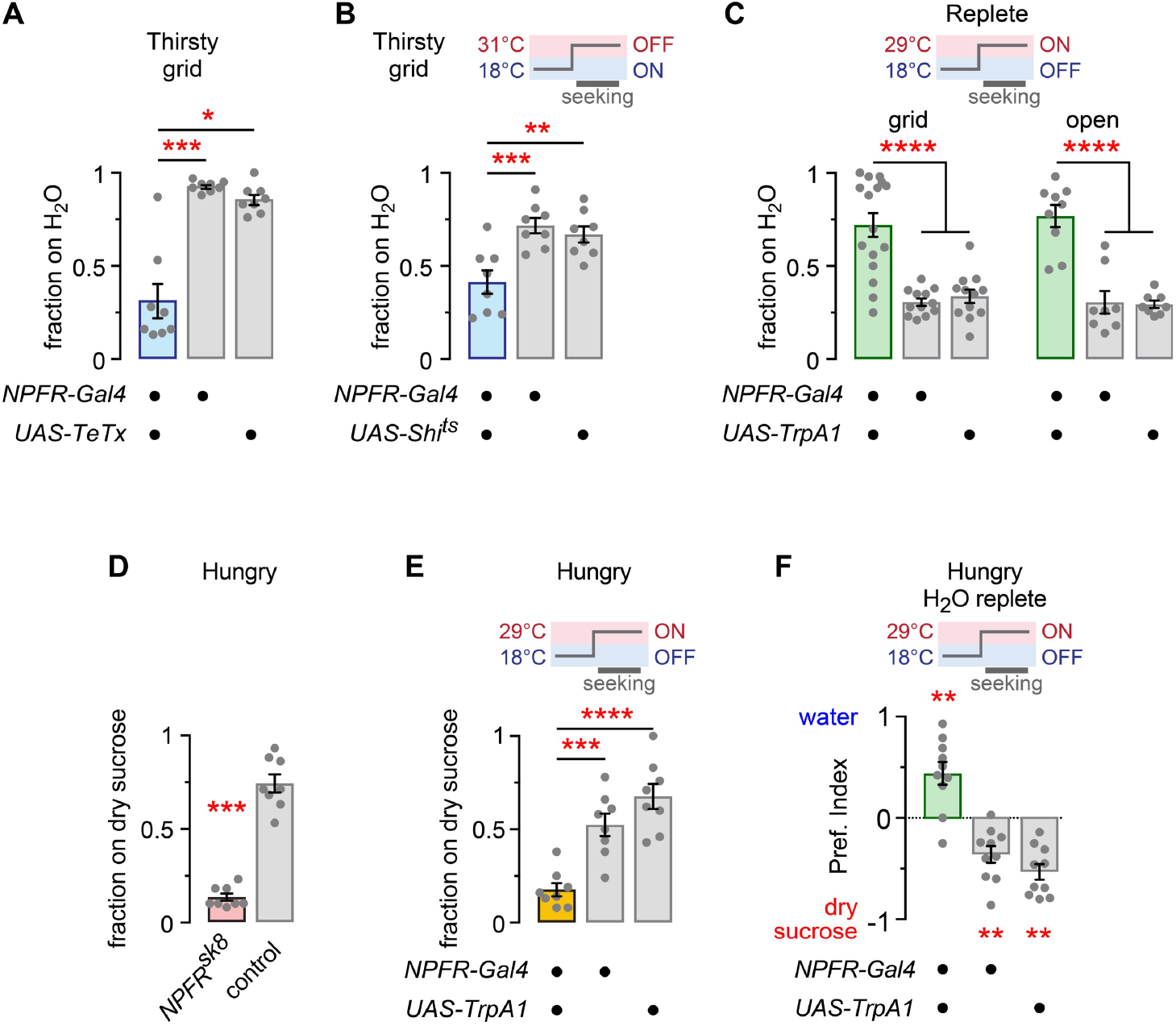
NPFR neuron activity is sufficient to promote water seeking and to regulate feeding behavior. **A**. NPFR neuron inactivation with the tetanus toxin light chain (*UAS-TeTx*) blocks thirsty water seeking. Kruskal-Wallis ANOVA with Dunn’s multiple comparisons test; *Gal4/UAS* vs. *Gal4, p* = 0.0001; *Gal4/UAS* vs. *UAS, p* = 0.0427. **B**. Acute synaptic silencing (*UAS-Shi*^*ts*^) of NPFR neurons decreases thirsty water seeking. One-way ANOVA with Holm-Šídák’s multiple comparisons test; *Gal4/UAS* vs. *Gal4, p* = 0.0007; *Gal4/UAS* vs. *UAS, p* = 0.0016. **C**. Acute depolarization (*UAS-TrpA1*) of NPFR neurons increases water seeking in water replete flies, with a gridded or an open water source. Left: Brown-Forsythe ANOVA with Dunnett’s T3 multiple comparisons test; *Gal4/UAS* vs. *Gal4, p* < 0.0001; *Gal4/UAS* vs. *UAS, p* < 0.0001. Right: One-way ANOVA with Holm-Šídák’s multiple comparisons test; *Gal4/UAS* vs. *Gal4, p* < 0.0001; *Gal4/UAS* vs. *UAS, p* < 0.0001. **D**. *NPFR*^*sk8*^ mutant flies fail to forage for food when wet starved. Mann-Whitney test, two-tailed, *p* = 0.0002. **E**. Acute depolarization (*UAS-TrpA1*) of NPFR neurons in wet starved flies decreases foraging for food. One-way ANOVA with Holm-Šídák’s multiple comparisons test; *Gal4/UAS* vs. *Gal4, p* = 0.0003; *Gal4/UAS* vs. *UAS, p* < 0.0001. **F**. Hungry but water replete (wet starved) flies given a choice between dry sucrose and water choose water when NPFR neurons are activated. Wilcoxon signed rank test. *Gal4*/*UAS, p* = 0.0078; *Gal4, p* = 0.0039; *UAS, p* = 0.0020.

### A subset of NPFR neurons promotes thirsty water seeking

To begin to identify the specific NPFR neurons that control thirsty seeking, we reduced NPFR expression in neurons that are labeled in four different *NPFR* genomic *enhancer-Gal4s* (**Figure 3A**). Three of the four, *R61H06-Gal4, R60G05-Gal4*, and *R65C12-Gal4*, resulted in reduced thirsty water seeking when driving *NPFR* RNAi (**Figure 3B**). Previously available whole brain immunohistochemical analysis revealed that each of these *enhancer-Gal4s* labeled complex patterns of neurons in the central brain (Jenett et al., 2012). We were unable to visually identify neurons that were common between all three *enhancer-Gal4s*. To narrow the potential site of action for NPFR, we employed *split-Gal4* technology, where the activation domain of *Gal4* was driven by the *R61H06* enhancer fragment, and the DNA-binding domain of *Gal4* was driven by the *R65C12* enhancer fragment. Reagents to generate *split-Gal4s* with *R60G05* were not available. Expression of either *NPFR* RNAi or TeTx with *R61H06:R65C12-spGal4* resulted in reduced thirsty water seeking (**Figure 3C**). Thus, NPFR water seeking neurons exist in the *R61H06:R65C12-spGal4* pattern. This *spGal4* was expressed in 5-7 pairs of neurons in the central brain and 10-15 pairs of neurons in the ventral nerve cord (**Figure 3D**). To better identify the neurons in the central brain, we used the multicolor flip out (MCFO) technique to stochastically label neuron subsets (**Figure 3E-I**). This technique unambiguously identified the presence of the two NPF-positive NPFR neurons, the L1-l and the P1 neurons (**Figure 3E,F**). The MCFO brains also revealed a dorsolateral neuron that is morphologically similar to SLP374 in the full adult female brain (FAFB) electron microscopy reconstruction and connectome (Dorkenwald et al., 2024; Schlegel et al., 2024) (**Figure 3G**). We also recovered ascending neurons that appear to be either AN_multi_124 or AN_multi_125 in the FAFB (**Figure 3H**). Two neurons were present in the *spGal4* but were not recovered in the MCFO screen. First, 1-2 neurons in the pars intercerebralis were labeled (**Figure 3D**). Second, a ventral lateral neuron was tentatively identified it as DNg30, based on the position and large size of its cell body and its unique axonal projections.

**Figure 3.**
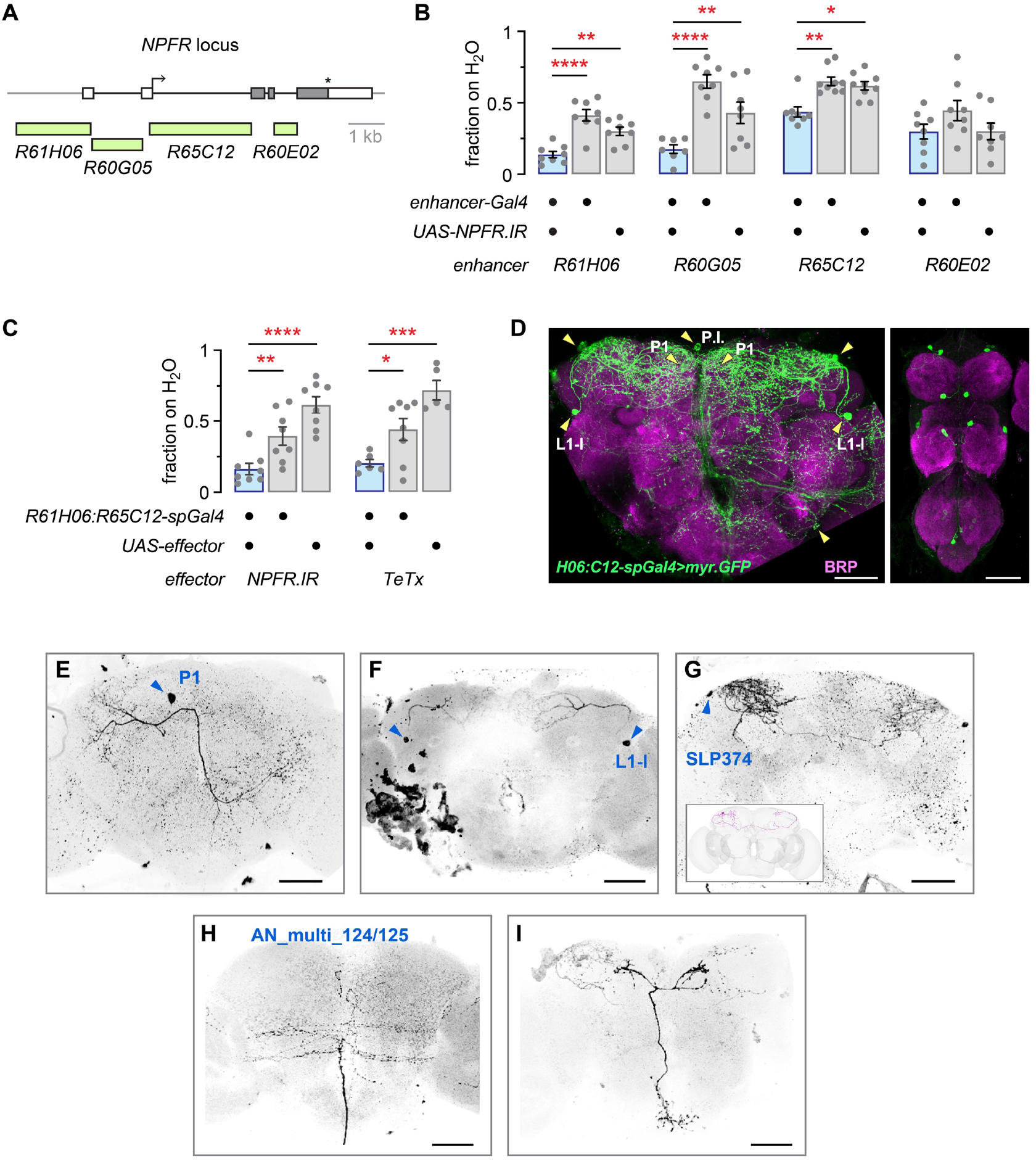
A subset of NPFR neurons promote thirsty water seeking. **A**. Genomic location of the DNA fragments used to create NPFR *enhancer-Gal4s*. **B**. Effect of reducing NPFR expression in the NPFR *enhancer-Gal4s*. Left: one-way ANOVA with Holm-Šídák’s multiple comparisons test; *Gal4/UAS* vs. *Gal4, p* < 0.0001; *Gal4/UAS* vs. *UAS, p* = 0.0016. Center left: One-way ANOVA with Holm-Šídák’s multiple comparisons test; *Gal4/UAS* vs. *Gal4, p* < 0.0001; *Gal4/UAS* vs. *UAS, p* = 0.0046. Center right: Kruskal-Wallis with Dunn’s multiple comparisons test; *Gal4/UAS* vs. *Gal4, p* = 0.0053; *Gal4/UAS* vs. *UAS, p* = 0.0143. Right: One-way ANOVA, *p* = 0.1677. **C**. Effect of *NPFR RNAi* (left) and *TeTx* (right) on thirsty water seeking when expressed in the *R61H06-Gal4*.*AD*;*R65C12-Gal4*.*DBD split-Gal4*. Left: one-way ANOVA with Holm-Šídák’s multiple comparisons test; *spGal4/UAS* vs. *spGal4, p* = 0.0077; *spGal4/UAS* vs. *UAS, p* < 0.0001. Right: one-way ANOVA with Holm-Šídák’s multiple comparisons test; *spGal4/UAS* vs. *spGal4, p* = 0.0180; *spGal4/UAS* vs. *UAS, p* = 0.0002. **D**. Expression pattern of *R61H06*:*R65C12*-*spGal4* in the central brain (left) and ventral nerve cord (right). Arrowheads point to cell bodies. Scale bars: 50 μm. **E**-**I**. Multicolor flip-out (MCFO) stochastic labeling reveals morphology of individual neurons in *R61H06*:*R65C12*-*spGal4*. Inset in **G** is the morphology of SLP374 in the FAFB. Arrowheads point to cell bodies. Scale bars: 50 μm.

### NPFR-positive neurons in the NPFR split-Gal4

We immunostained *R61H06:R65C12-spGal4* expressing myristoylated GFP with antibodies to GFP and NPFR to help reveal which neurons in the *spGal4* are NPFR-positive (**Figure 4**). As expected, the L1-l and P1 neurons were NPFR-positive (**Figure 4A,B**). Also NPFR-positive was the neuron in the pars intercerebralis (**Figure 4C,D**). We were unable to detect NPFR in either of the other two central brain neurons, the putative SLP374 neuron and the putative DNg30 neuron (**Figure 4D,E**). In the ventral nerve cord two pairs of as-yet unidentified neurons were weakly positive for NPFR (**Figure 4F**).

**Figure 4.**
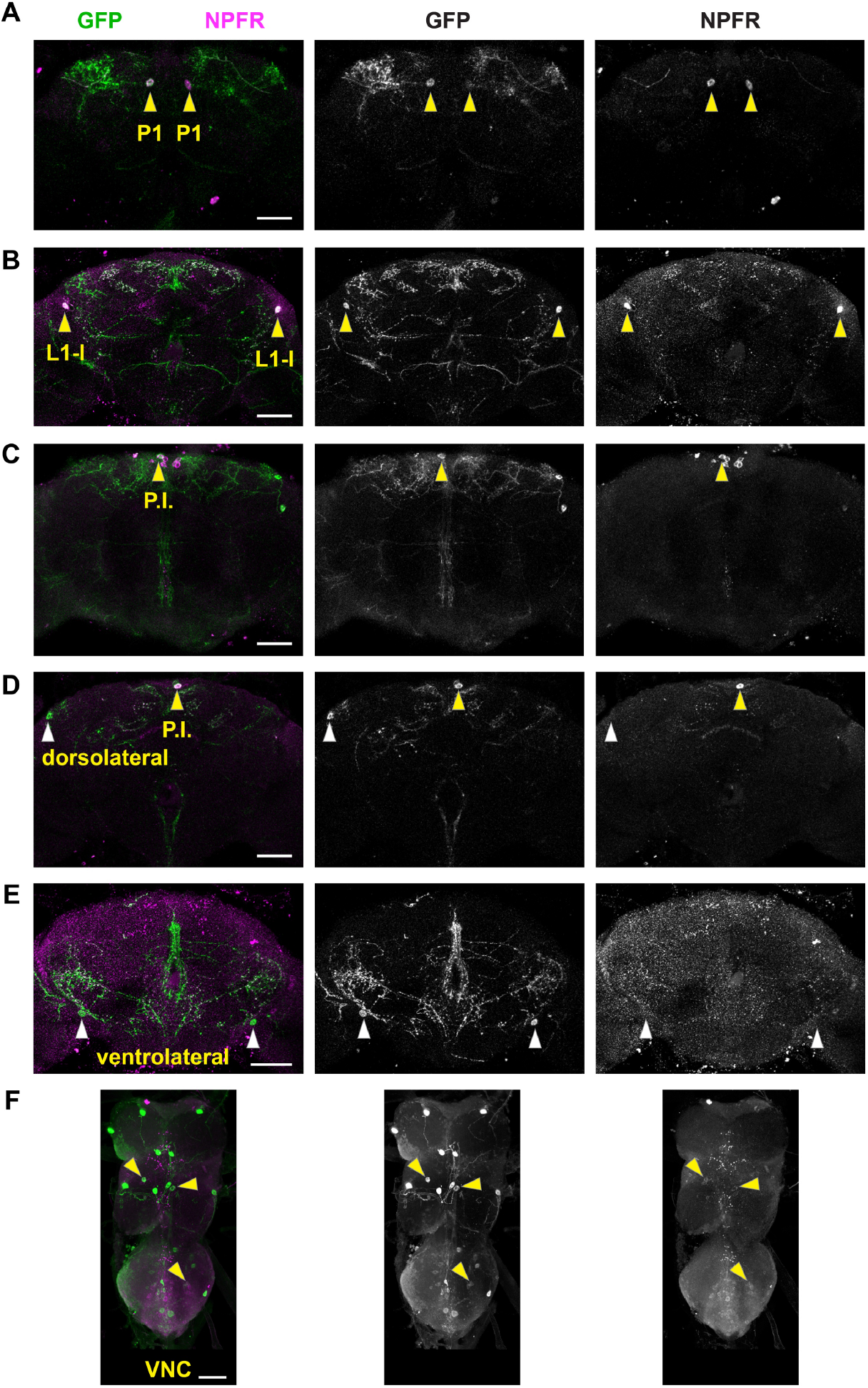
Immunostaining of *R61H06*:*R65C12*-*spGal4>UAS-myr*.*GFP* central brains (**A**-**E**) and ventral nerve cords (**F**) with antibodies to GFP (green) and NPFR (magenta). Each panel is a maximum intensity projection of a z-series to reveal individual cell bodies. Arrowheads point to positions of GFP-positive cell bodies that are positive (yellow) or negative (white) for NPFR. Scale bars: 50 μm.

### NPFR acts in the L1-l neuron to promote thirsty water seeking

Water deprivation increases neuronal activity in NPFR-positive L1-l and P1 neurons (Landayan et al., 2021). Neurons in the neuroendocrine pars intercerebralis regulate water ingestion, suggesting that they may also be a site of NPFR action for thirsty water seeking (González Segarra et al., 2023). We created new *spGal4s* to test the water seeking role of the L1-l and the P1 neurons. To do this, we used an *NPF-DBD* hemidriver and screened for *-AD* hemidrivers that together labeled small numbers or individual neurons. We identified *R20A02-AD;NPF-DBD* as an L1-l-specific driver, and *R23E12-AD;NPF-DBD* as a P1-specific *spGal4* driver (**Figure 5A**,**B**). Expression of NPFR RNAi in the L1-l neuron reduced thirsty water seeking, whereas it caused a trend towards reduced water seeking in the P1 neuron (**Figure 5A’,B’**). Most or all NPFR-positive neurons in the pars intercerebralis are also positive for the neuropeptide DH44, that we confirmed (**Figure 5C**) (Ryvkin et al., 2024). Thus, we reduced NPFR expression in DH44 neurons with *Dh44-Gal4*, however we did not observe an effect on thirsty water seeking (**Figure 5C’**). Taken together, these results indicate that the L1-l neuron promotes water seeking by receiving NPF input through the NPFR receptor. Because the effect on thirsty water seeking of NPFR knockdown in the L1-l was moderate compared to *NPFR-Gal4*, we suspect that additional NPFR neurons also support thirsty water seeking.

**Figure 5.**
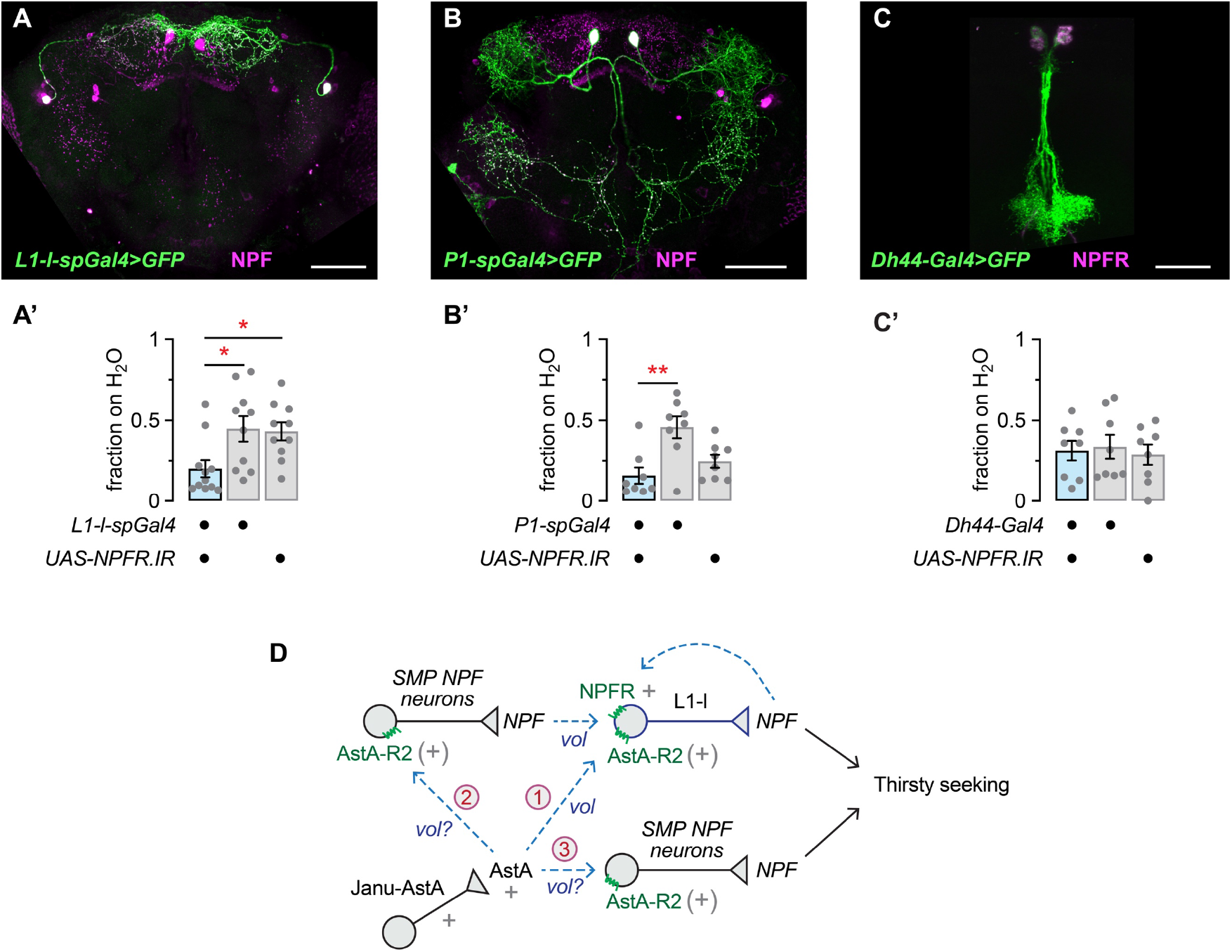
Effects on thirsty water seeking of reducing NPFR expression in individual NPFR neurons. **A, A’**. *L1-l-spGal4* (*R20A02-Gal4*.*AD*;*NPF-Gal4*.*DBD*) neurons immunostained for GFP and NPF (**A**). Reduced thirsty water seeking with reduced NPFR expression in *L1-l-spGal4* neurons (**A’**). One-way ANOVA with Holm-Šídák’s multiple comparisons test; *spGal4/UAS* vs. *spGal4, p* = 0.0177; *spGal4/UAS* vs. *UAS, p* = 0.0269. **B, B’**. *P1-spGal4* (*R23E12-Gal4*.*AD*;*NPF-Gal4*.*DBD*) neurons immunostained for GFP and NPF (**B**). Non-significant trend towards decreased thirsty water seeking with reduced NPFR expression in *P1-spGal4* neurons (**B’**). One-way ANOVA with Holm-Šídák’s multiple comparisons test; *spGal4/UAS* vs. *spGal4, p* = 0.0015; *spGal4/UAS* vs. *UAS, p* = 0.2554. **C, C’**. *DH44-Gal4* neurons immunostained for GFP and NPFR (**C**). No effect on thirsty water seeking with reduced NPFR expression in *Dh44-Gal4* neurons (**C’**). One-way ANOVA, *p* = 0.8742. **D**. Circuit models. “+” indicates characterized role, “(+)” for AstA-R2 functioning on NPF neurons, dashed arrows and “vol” indicate proposed volumetric transmission, and circled numbers designate the proposed models described in the Discussion. Scale bars: 50 μm.

## Discussion

We identified NPFR, the receptor for the neuropeptide NPF, and neurons that express NPFR as critical for the motivated ingestive behaviors water seeking, water intake, and feeding. Using intersectional genetic techniques, we identified the bilaterally symmetric and NPF-positive L1-l neuron as requiring NPFR for the promotion of thirsty water seeking. Here we discuss the position of NPFR function in the emerging circuitry for innate thirst behavior in the Drosophila brain, and how the L1-l neuron may integrate into brain circuits that regulate memories and the valuation of sensory input.

In mammals, the NPF ortholog NPY and its receptors are studied mainly for their role in suppressing hunger and modulating memory, however there are a few reports that implicate NPY and the Y2 and Y4 NPY receptors in thirst (Morley and Flood, 1989; Wultsch et al., 2006). Hunger and thirst both strongly drive seeking and ingestion, and they interact behaviorally and at the neural circuit level in rodents (Augustine et al., 2020; Encarnacion-Rivera et al., 2025). Individual neurons in the Drosophila brain can regulate both hunger and thirst (Jourjine, 2017; Landayan et al., 2021; Shiu et al., 2022; González Segarra et al., 2023). NPF itself oppositely regulates hunger and thirst in flies (Landayan et al., 2021).

A Janu-AstA to NPF to NPFR circuit is supported by prior evidence: NPF functions downstream of Janu-AstA neurons via the AstA receptor AstA-R2 to regulate thirsty water seeking, and the L1-l neurons are activated by thirst (Landayan et al., 2021). Neither the available connectome reconstructions of the Drosophila brain nor anterograde synaptic tracing experiments support the L1-l as being directly postsynaptic to Janu-AstA (not shown). Given these biological constraints, the following thirst circuits are possible (**Figure 5D**). 1) AstA from Janu-AstA may function through volumetric transmission, also known as extra-synaptic diffusion, to reach NPFR expressing L1-l neurons within the superior medial protocerebrum (SMP) (Girven et al., 2022). Janu-AstA elaborates presynaptic release sites in the SMP, and L1-l neuronal processes pass through and near the Janu-AstA release zone. NPFR on the L1-l may receive autocrine NPF directly from the L1-l: AstA released from Janu-AstA neurons likely regulates the release of NPF from L1-l neurons. 2) An unidentified NPF neuron may function upstream of NPFR on the L1-l neurons, acting between Janu-AstA and L1-l. Three neurons synaptically connect Janu-AstA to the L1-l: MBON35 (receiving 24 or 3.8% of Janu-AstA synapses, and delivering 75 or 4.2% of all synapses to the L1-l in the hemibrain), OviIN (receiving 28 or 4.5% of Janu-AstA synapses, and delivering 34 or 1.9% synapses to the L1-l), and SMP081 (two per hemisphere, receiving 31-33 or 4.9-5.3% of Janu-AstA synapses, and delivering 18-36 or 1-1.9% synapses to the L1-l). All three intermediate neuron types are considered “rich-club” neurons that make many synaptic connections across multiple brain regions. For example, MBON35 receives 21,335 synaptic inputs and delivers 1746 synaptic outputs in the hemibrain. Moreover, none of these bridging neurons is likely to be NPF-positive, based on prior morphological characterization of the NPF neurons. Hence, in the second model volume transmission is likely necessary to allow for one of the other NPF neurons to bridge between Janu-AstA and L1-l. Candidate neurons based on their innervation of the SMP include the male-specific NPF-M neuron and the NPF-positive neurons in the evening groups of the circadian clock, the LNd2, LNd3, LNd6, and 5th LNv (Lee et al., 2006; Kim et al., 2013; Liu et al., 2019; Shafer et al., 2022). The small DMs are a final group of NPF neurons that arborize in the medial part of the dorsal protocerebrum, however their identity in the connectome is not yet known. The small DMs encode positive valence (Shao et al., 2017). 3) Janu-AstA and L1-l may function in parallel circuits to promote thirsty water seeking.

What role might the L1-l play in thirsty seeking? The identity and function of L1-l output neurons can provide clues. In the FAFB there are only 8 types of output neurons, with the left L1-l making a total of 37 synapses to postsynaptic neurons and the right 47. The two outputs receiving the most synapses from the L1-l are MBON35 (10 synapses per L1-l, FAFB) and SMP108 (9-12 synapses per L1-l, FAFB). Although a substantial number of L1-l output synapses are dedicated to the MBON35 and SMP108 (19-22 out of 50-55 total), the L1-ls represent only 0.12-0.15% of MBON35 or SMP108 inputs. Interestingly, all L1-l outputs are classified as “rich-club”. We suggest three possible interpretations. First, L1-ls signal a bias to “rich-club” neurons to generally tune or sensitize a brain state. For example, coincident activity of neurons from separate circuits onto a “rich-club” neuron may alter its likelihood of firing. This type of encoding could be part of internal state encoding where the influence of the state is graded over time to alter the probability of the execution of a thirst related behavior such as increased locomotion or sensory tuning. Second, the main targets of the L1-l may be due to volumetric transmission, rather than direct synaptic connections. Finally, it remains possible that existing connectomes do not capture important L1-l outputs.

The L1-l neuron is also named the DAL2 (Dorsal Anterior Lateral) neuron, and it regulates long-term memory consolidation through their action on the PPL1 aversive dopamine neurons (Feng et al., 2021). Specifically, L1-l neurons inhibit the PPL1 neurons to decrease their interference with long term memory consolidation. Despite no direct synaptic connectivity, L1-l release of NPF is critical for the repression of PPL1 neuron electrical activity through the NPFR receptor during memory consolidation. By analogy, elevated L1-l activity in the thirsty state may cause repression of PPL1 neurons, leading to changes in the sensitivity of the mushroom body circuitry to sensory inputs and positive valence dopaminergic inputs. Thirst permits appetitive long term memory formation for water reward that is thought to involve specific sets of positive valence PAM dopaminergic neurons that innervate the mushroom body (Lin et al., 2024). A subset of positive valence PAM neurons are activated by water ingestion in thirsty flies and they regulate types of water-related memory formation (Lee et al., 2025). Similarly, foraging for food when hungry is regulated by NPFR on PPL1 neurons, suggesting that NPF-to-dopamine neuron signaling regulates a valence aspect of seeking related to ingestive behavior (Tsao et al., 2018). Finally, the L1-l output neuron SMP108 is presynaptic to the PAM reward dopamine neurons, and it functions in a type of learning termed second-order conditioning (Yamada et al., 2023).

## Acknowledgements

This research was supported by startup funds from the University of California, Merced. We thank Michael Texada for providing us with the *NPFR-Gal4* transgene prior to its publication, and for his insights into neuromodulators in Drosophila.

## Notes

**Conflicts of Interest** Authors report no conflict of interest.

### Competing Interest Statement

The authors have declared no competing interest.

## References

Ameku T, Yoshinari Y, Texada MJ, Kondo S, Amezawa K, Yoshizaki G, Shimada-Niwa Y, Niwa R (2018) Midgut-derived neuropeptide F controls germline stem cell proliferation in a mating-dependent manner. PLoS Biol 16:e2005004.

Augustine V, Lee S, Oka Y (2020) Neural Control and Modulation of Thirst, Sodium Appetite, and Hunger. Cell 180:25–32.

Beshel J, Zhong Y (2013) Graded Encoding of Food Odor Value in the Drosophila Brain. J Neurosci 33:15693–15704.

Betley JN, Xu S, Cao ZFH, Gong R, Magnus CJ, Yu Y, Sternson SM (2015) Neurons for hunger and thirst transmit a negative-valence teaching signal. Nature 521:180–185.

Chu L-A, Tai C-Y, Chiang A-S (2024) Thirst-driven hygrosensory suppression promotes water seeking in Drosophila. Proc Natl Acad Sci U S A 121:e2404454121.

Dethier VG, Evans DR (1961) The Physiological Control of Water Ingestion in the Blowfly. Biol Bull Mar Biol Lab 121:108–116.

Dorkenwald S et al. (2024) Neuronal wiring diagram of an adult brain. Nature 634:124–138.

Encarnacion-Rivera L, Deisseroth K, Luo L (2025) Neurobiology of Thirst and Hunger Drives. Annu Rev Neurosci.

Enjin A, Zaharieva EE, Frank DD, Mansourian S, Suh GSB, Gallio M, Stensmyr MC (2016) Humidity Sensing in Drosophila. Curr Biol 26:1352–1358.

Feng K-L, Weng J-Y, Chen C-C, Abubaker MB, Lin H-W, Charng C-C, Lo C-C, de Belle JS, Tully T, Lien C-C, Chiang A-S (2021) Neuropeptide F inhibits dopamine neuron interference of long-term memory consolidation in Drosophila. iScience 24:103506.

Fitzsimons JT (1972) Thirst. Physiol Rev 52:468–561.

Frank DD, Enjin A, Jouandet GC, Zaharieva EE, Para A, Stensmyr MC, Gallio M (2017) Early Integration of Temperature and Humidity Stimuli in the Drosophila Brain. Curr Biol 27:2381-2388.e4.

Gáliková M, Dircksen H, Nässel DR (2018) The thirsty fly: Ion transport peptide (ITP) is a novel endocrine regulator of water homeostasis in Drosophila. PLoS Genet 14:e1007618.

Gao J, Zhang S, Deng P, Wu Z, Lemaitre B, Zhai Z, Guo Z (2024) Dietary L-Glu sensing by enteroendocrine cells adjusts food intake via modulating gut PYY/NPF secretion. Nat Commun 15:3514.

Girven KS, Mangieri L, Bruchas MR (2022) Emerging approaches for decoding neuropeptide transmission. Trends Neurosci 45:899–912.

Gizowski C, Bourque CW (2018) The neural basis of homeostatic and anticipatory thirst. Nat Rev Nephrol 14:11–25.

Gizowski C, Zaelzer C, Bourque CW (2016) Clock-driven vasopressin neurotransmission mediates anticipatory thirst prior to sleep. Nature 537:685–688.

González Segarra AJ, Pontes G, Jourjine N, Del Toro A, Scott K (2023) Hunger- and thirst-sensing neurons modulate a neuroendocrine network to coordinate sugar and water ingestion. Elife 12:RP88143.

Jenett A et al. (2012) A GAL4-driver line resource for Drosophila neurobiology. Cell Rep 2:991–1001.

Ji F, Zhu Y (2015) A novel assay reveals hygrotactic behavior in Drosophila. PLoS ONE 10:e0119162.

Jourjine N (2017) Hunger and thirst interact to regulate ingestive behavior in flies and mammals. Bioessays 39.

Kim WJ, Jan LY, Jan YN (2013) A PDF/NPF Neuropeptide Signaling Circuitry of Male Drosophila melanogaster Controls Rival-Induced Prolonged Mating. Neuron 80:1190–1205.

Knecht ZA, Silbering AF, Cruz J, Yang L, Croset V, Benton R, Garrity PA (2017) Ionotropic Receptor-dependent moist and dry cells control hygrosensation in Drosophila. Elife 6.

Krashes MJ, DasGupta S, Vreede A, White B, Armstrong JD, Waddell S (2009) A neural circuit mechanism integrating motivational state with memory expression in Drosophila. Cell 139:416–427.

Landayan D, Wang BP, Zhou J, Wolf FW (2021) Thirst interneurons that promote water seeking and limit feeding behavior in Drosophila. Elife 10:e66286.

Lee G, Bahn JH, Park JH (2006) Sex- and clock-controlled expression of the neuropeptide F gene in Drosophila. PNAS 103:12580–12585.

Lee W-P, Chiang M-H, Chao Y-P, Wang Y-F, Chen Y-L, Lin Y-C, Jenq S-Y, Lu J-W, Fu T-F, Liang J-Y, Yang K-C, Chang L-Y, Wu T, Wu C-L (2025) Dynamics of two distinct memory interactions during water seeking in Drosophila. Proc Natl Acad Sci U S A 122:e2422028122.

Limbania D, Turner GL, Wasserman SM (2023) Dehydrated Drosophila melanogaster track a water plume in tethered flight. iScience 26:106266.

Lin S, Owald D, Chandra V, Talbot C, Huetteroth W, Waddell S (2014) Neural correlates of water reward in thirsty Drosophila. Nat Neurosci 17:1536–1542.

Lin Y-C, Wu T, Wu C-L (2024) The Neural Correlations of Olfactory Associative Reward Memories in Drosophila. Cells 13:1716.

Liu W, Ganguly A, Huang J, Wang Y, Ni JD, Gurav AS, Aguilar MA, Montell C (2019) Neuropeptide F regulates courtship in Drosophila through a male-specific neuronal circuit. Elife 8.

Marin EC, Büld L, Theiss M, Sarkissian T, Roberts RJV, Turnbull R, Tamimi IFM, Pleijzier MW, Laursen WJ, Drummond N, Schlegel P, Bates AS, Li F, Landgraf M, Costa M, Bock DD, Garrity PA, Jefferis GSXE (2020) Connectomics Analysis Reveals First-, Second-, and Third-Order Thermosensory and Hygrosensory Neurons in the Adult Drosophila Brain. Curr Biol 30:3167-3182.e4.

Morley JE, Flood JF (1989) The effect of neuropeptide Y on drinking in mice. Brain Res 494:129–137.

Ryvkin J, Bentzur A, Shmueli A, Tannenbaum M, Shallom O, Dokarker S, Benichou JIC, Levi M, Shohat-Ophir G (2021) Transcriptome Analysis of NPFR Neurons Reveals a Connection Between Proteome Diversity and Social Behavior. Front Behav Neurosci 15:628662.

Ryvkin J, Omesi L, Kim Y-K, Levi M, Pozeilov H, Barak-Buchris L, Agranovich B, Abramovich I, Gottlieb E, Jacob A, Nässel DR, Heberlein U, Shohat-Ophir G (2024) Failure to mate enhances investment in behaviors that may promote mating reward and impairs the ability to cope with stressors via a subpopulation of Neuropeptide F receptor neurons. PLoS Genet 20:e1011054.

Schlegel P et al. (2024) Whole-brain annotation and multi-connectome cell typing of Drosophila. Nature 634:139–152.

Shafer OT, Gutierrez GJ, Li K, Mildenhall A, Spira D, Marty J, Lazar AA, Fernandez M de la P (2022) Connectomic analysis of the Drosophila lateral neuron clock cells reveals the synaptic basis of functional pacemaker classes. Elife 11:e79139.

Shao L, Saver M, Chung P, Ren Q, Lee T, Kent CF, Heberlein U (2017) Dissection of the Drosophila neuropeptide F circuit using a high-throughput two-choice assay. Proc Natl Acad Sci USA 114:E8091–E8099.

Shiu PK, Sterne GR, Engert S, Dickson BJ, Scott K (2022) Taste quality and hunger interactions in a feeding sensorimotor circuit. Elife 11:e79887.

Tsao C-H, Chen C-C, Lin C-H, Yang H-Y, Lin S (2018) Drosophila mushroom bodies integrate hunger and satiety signals to control innate food-seeking behavior. Elife 7.

Wen T, Parrish CA, Xu D, Wu Q, Shen P (2005) Drosophila neuropeptide F and its receptor, NPFR1, define a signaling pathway that acutely modulates alcohol sensitivity. Proc Natl Acad Sci U S A 102:2141–2146.

Wu Q, Zhao Z, Shen P (2005) Regulation of aversion to noxious food by Drosophila neuropeptide Y- and insulin-like systems. Nat Neurosci 8:1350–1355.

Wultsch T, Painsipp E, Donner S, Sperk G, Herzog H, Peskar BA, Holzer P (2006) Selective increase of dark phase water intake in neuropeptide-Y Y2 and Y4 receptor knockout mice. Behav Brain Res 168:255–260.

Xu J, Li M, Shen P (2010) A G-protein-coupled neuropeptide Y-like receptor suppresses behavioral and sensory response to multiple stressful stimuli in Drosophila. J Neurosci 30:2504–2512.

Yamada D, Bushey D, Li F, Hibbard KL, Sammons M, Funke J, Litwin-Kumar A, Hige T, Aso Y (2023) Hierarchical architecture of dopaminergic circuits enables second-order conditioning in Drosophila. Elife 12:e79042.

Yoshinari Y, Kosakamoto H, Kamiyama T, Hoshino R, Matsuoka R, Kondo S, Tanimoto H, Nakamura A, Obata F, Niwa R (2021) The sugar-responsive enteroendocrine neuropeptide F regulates lipid metabolism through glucagon-like and insulin-like hormones in Drosophila melanogaster. Nat Commun 12:4818.

Yuan X, Li H, Guo F (2024) Temperature cues are integrated in a flexible circadian neuropeptidergic feedback circuit to remodel sleep-wake patterns in flies. PLoS Biol 22:e3002918.

Zandawala M, Nguyen T, Balanyà Segura M, Johard HAD, Amcoff M, Wegener C, Paluzzi J-P, Nässel DR (2021) A neuroendocrine pathway modulating osmotic stress in Drosophila. PLoS Genet 17:e1009425.

